# Cell Type Architecture and Positional Gene Gradients in an Adult Animal at Subcellular Resolution

**DOI:** 10.64898/2026.02.19.705280

**Authors:** Maoqin Sun, Yuxiaofei Wang, Kai Han, Lidong Guo, Yue Chen, Yao Li, Yaru Lin, Xiawei Liu, Zhi Huang, Qun Liu, Wenjie Guo, Rui Zhang, Wandong Zhao, Langchao Liang, Xiaoyu Wei, Li Zhou, Xuebin Mao, Jiaqi Wang, Weijian Wu, Hongwei Pan, Tao Yang, He Zhang, Xiaoshan Su, Shanshan Liu, Wenwei Zhang, Longqi Liu, Søren Tvorup Christensen, Jifeng Fei, Xin Liu, Ying Gu, Jian Wang, Huanming Yang, Gang Pei, Guangyi Fan, Xun Xu, Hanbo Li, Mengyang Xu, An Zeng

## Abstract

The spatial organization of cell types and gene expression underlies tissue architecture and organismal physiology. However, resolving complete cellular and molecular landscapes within a three-dimensional context remains challenging, particularly in organisms with high cellular heterogeneity and complex geometry. Here, we developed a semi-supervised workflow to reconstruct high-resolution three-dimensional spatial transcriptomic models of the planarian *Schmidtea mediterranea* at single-cell resolution. Planarians are basal bilaterians capable of regenerating body structures after injury. Our reconstruction maps the distribution of cell types and spatially patterned gene expression across the intact organism and identifies 119 candidate positional control genes (PCGs). These genes are expressed across muscle, neural, and epidermal populations. Functional perturbation of selected candidates, including *pitx3*, *ptpn11*, *pi4ka*, *upf3b*, and *cul1*, revealed roles in regeneration following injury. In addition, we identify intestinal cells as prominent components of the microenvironment surrounding adult *tgs1*⁺ pluripotent stem cells (neoblasts). Together, these results demonstrate the utility of three-dimensional spatial transcriptomic reconstruction for mapping cellular architecture and positional gene regulation in complex adult organisms.

## Introduction

The spatial organization of cell types and gene expression underlies tissue architecture and organismal physiology. In multicellular organisms, positional information coordinates cell identity, tissue patterning, and regeneration [1], yet how this information is organized and maintained across an intact three-dimensional body remains incompletely understood. Defining cell type architecture and positional gene gradients at organismal scale is therefore essential for understanding how complex body plans are established and repaired.

Planarians, renowned for their regenerative capacity, provide a valuable model for investigating stem cell-mediated tissue repair and regeneration [2, 3]. In these organisms, adult pluripotent stem cells, or neoblasts, are essential for regenerating lost tissues [4, 5]. Stem cell behaviors, including self-renewal, differentiation, and migration, are regulated by the surrounding microenvironment, comprising local cellular and molecular interactions [6, 7]. Although substantial progress has been made in defining the molecular properties of planarian stem cells, the spatial organization and influence of the microenvironment on neoblast behavior remain incompletely characterized [8].

Regeneration in planarians is guided by positional cues that regulate body axis formation and tissue patterning. These cues involve genes termed positional control genes (PCGs) [9], which exhibit region-specific expression and are usually linked to RNAi-induced patterning defects [10–12]. PCGs have been primarily described as being expressed in muscle; however, whether their expression is restricted to muscle, how positional signals are transmitted to stem cells to re-establish body axes, and what molecular cascades ultimately restore functional three-dimensional tissue architecture remain incompletely understood [13, 14]. Addressing these questions requires complete profiling organisms, cells, and genes across multiple spatial scales in three dimensions.

Recent advances in spatial transcriptomics have enabled in situ visualization of gene expression within tissues [15–17]. However, most approaches remain limited to two-dimensional sections or to resolutions insufficient to fully resolve small cells such as neoblasts (5–10 μm in diameter). Planarians also present anatomical challenges, including a compact dorsoventral (D/V) axis and substantial heterogeneity along the anteroposterior (A/P) and mediolateral (M/L) axes [18]. Two-dimensional sections, while informative, do not fully capture the organism-wide spatial context of stem cell niches.

To address these challenges and obtain a comprehensive view of spatial gene expression and tissue organization, a high-resolution three-dimensional spatial transcriptomic model capable of resolving individual cells and their neighboring components is required. However, generating accurate 3D reconstructions of flexible and anatomically complex organisms such as planarians remains technically demanding, particularly when relying solely on automated algorithms [19, 20]. Major obstacles include precise segmentation of small tissue structures and individual cells in the presence of image noise and variability, as well as reliable alignment of serial sections to produce a coherent three-dimensional model. These challenges are exacerbated in soft-bodied organisms with intricate anatomical features.

Here, we develop a semi-supervised reconstruction strategy optimized for high-resolution spatial transcriptomics using the Stereo-seq platform [21]. This workflow integrates enhanced image processing, customized segmentation and alignment algorithms, and targeted manual curation to improve reconstruction accuracy and robustness. Applying this pipeline to adult asexual planarians, we cryosectioned homeostatic animals into 27 consecutive serial sections at 10 μm thickness and profiled each section at 715 nm resolution in the X–Y plane without interval loss. Using this approach, we reconstructed a whole-animal 3D model comprising 893,703 high-quality segmented cells, enabling systematic analysis of spatial gene expression patterns and tissue organization. Spatial proximity analysis further identified intestinal cells as prominent neighboring components of neoblasts within the broader tissue environment. Together, these results establish a scalable strategy for organism-scale three-dimensional spatial transcriptomic reconstruction and provide a foundation for examining cell-type architecture and positional gene gradients.

## Results

### An end-to-end semi-supervised framework for 3D spatial transcriptomic reconstruction

To overcome the limitations of single-section spatial transcriptomics and enable organism-scale three-dimensional analysis of cell types and gene expression, we established a semi-supervised reconstruction workflow using the planarian model. Planarians pose specific challenges for 3D reconstruction due to their complex cellular composition, dispersed pluripotent stem cells (neoblasts), regionally specialized cell types, and pronounced anterior–posterior to dorsal–ventral axis ratio. The soft and deformable body structure also complicates section alignment and accurate anatomical registration. To address these constraints, we integrated image registration, cell segmentation, and serial alignment algorithms into a unified semi-supervised framework optimized for small, flexible tissues and high-resolution spatial transcriptomics data (Fig. 1A), leveraging the Stereo-seq platform [21].

**Fig. 1.**
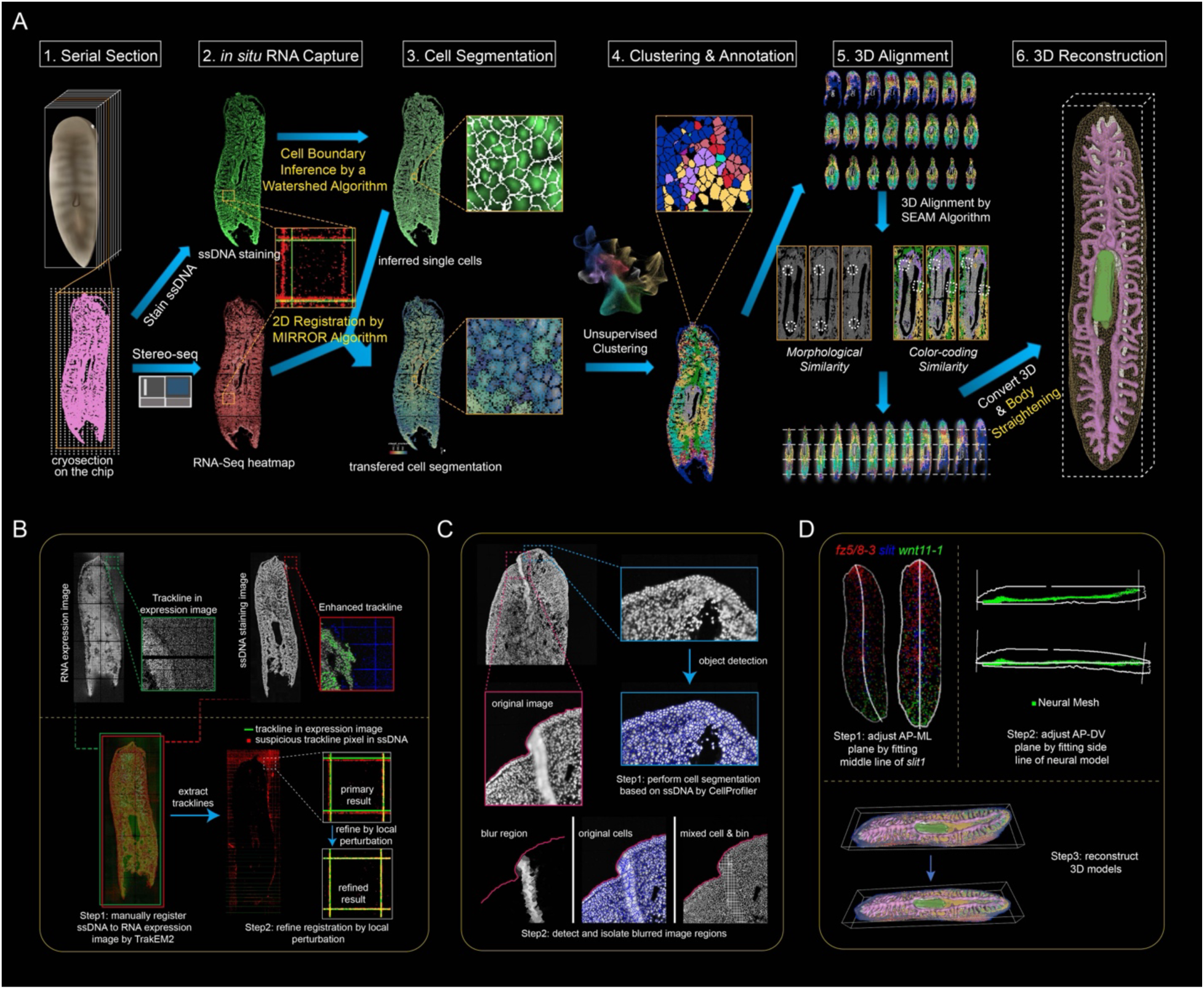
Workflow of 3D virtual reconstruction of planarian at single-cell resolution. (A) Schematic representation of the 3D virtual reconstruction process for the entire planarian. Key steps include sectioning, in situ barcoding, cell segmentation, clustering and annotation, alignment, and final 3D reconstruction. (B) Flowchart of the two-step 2D multimodal registration process using the MIRROR algorithm. Step 1: Rough alignment of ssDNA staining and color-coded gene expression images based on tissue morphological borders and track-line grids in TrakEM2. Step 2: Fine registration is achieved by an automated algorithm that matches track-line pairs and performs a global search to maximize pixel overlap. (C) Flowchart of the cell segmentation strategy. Step 1: Identification of nuclei locations in ssDNA staining images using the Otsu adaptive thresholding method, followed by cell area prediction via the Mutex Watershed algorithm in a customized CellProfiler pipeline. Step 2: Detection of complex or blurred regions using the Laplacian algorithm, with these regions replaced by a bin15 partition for improved accuracy. (D) Flowchart outlining the body straightening strategy. Step 1: Whole-body straightening in the A/P-M/L (x, y) plane based on anatomical centerline positioning determined by expression patterns of canonical A/P and M/L PCGs. Step 2: Straightening along the A/P-D/V (x, z) plane is achieved through the reconstruction of a neural mesh model with flattened topography. Universal transformations are then applied to the spatial coordinates of individual cells and reconstructed 3D models.

We first applied high-resolution Stereo-seq to profile individual slices of an adult asexual planarian at 715 nm resolution. The animal was cryosectioned along the dorsoventral (D/V) axis into 27 consecutive slices (fig. S1A), yielding a total of 64,479,815 spots across all sections (data S1). Next, we aligned and registered the single-stranded DNA (ssDNA) staining images with the spatial gene expression coordinates using the MIRROR (Multimodal Image Registration by tRackline On-chip with enhanced Resolution) pipeline (Materials and Methods; Fig. 1, A and B). This step enabled accurate multimodal spatial integration of tissue sections for downstream analysis.

To segment individual cells from the ssDNA-staining images, we applied the customized CellProfiler pipeline [22], which incorporated adaptive thresholding (Otsu method), contrast enhancement, and the Mutex Watershed algorithm. To mitigate segmentation errors caused by densely packed nuclei or out-of-focus regions, we implemented Laplacian blur detection to identify low-quality pixels, which were subsequently aggregated into bin-level partitions approximating planarian cell size (Fig. 1C). RNA capture spots were then assigned to segmented cell boundaries based on spatial overlap, generating cell-level gene expression matrices for subsequent analyses.

To ensure accurate spatial registration across the entire organism, we employed Seurat integrative analysis to match shared cell types between serial sections [23]. Differentially expressed genes were used to annotate clusters, which were further validated against established planarian gene databases [24, 25]. This provided a reliable framework for annotating cell types, which were color-coded and mapped onto their spatial coordinates, providing morphological features for subsequent 3D alignment.

For the iterative alignment of serial sections, we utilized the SEAM (Serial sEction Alignment by anatomical and MRNA expression similarity) pipeline (Materials and Methods). This alignment process involved the application of affine transformation matrices to integrate images, cells, and RNA spots into a cohesive coordinate system. Each point within this system was annotated with key attributes, including cell identity and gene expression values. To enhance visualization quality and quantify tissue volume, we constructed organ and body meshes based on the spatial distribution of each cell lineage within the planarian (Fig. 1A).

Because embedding during tissue preparation can introduce deformation, particularly along the anterior–posterior (A/P) axis, we computationally corrected body curvature in the reconstructed model. This correction was guided by body axis markers (*fz-5/8-3*, *wnt11-1*) and internal anatomical landmarks such as the ventral nerve cords, improving anatomical consistency across sections (Fig. 1D). Together, these components constitute WACCA (Whole-Animal reconstruction pipeline using 2D multimodal registration, Cell segmentation, Clustering, and Annotation), an end-to-end framework for constructing and analyzing three-dimensional models from high-resolution spatial transcriptomic data. This pipeline addresses alignment challenges associated with complex, deformable tissues and limited direct anatomical correspondence between adjacent sections.

### Whole-animal 3D reconstruction reveals complex cell-type organization

Using the WACCA workflow, we constructed a complete virtual representation of a planarian, revealing the diversity of cell types within the organism. By integrating images, RNA spots, and segmented cells into a unified coordinate system, we generated a 3D dataset that encapsulates the complexity of planarian biology (Fig. 2A). This dataset supports multi-scale analysis, including subcellular gene expression profiles, cell-type atlases, and tissue-level organization within organ and body models. In total, the reconstruction comprises 893,703 high-quality segmented cells, providing a detailed 3D transcriptomic model of an adult planarian (Fig. 2A).

**Fig. 2.**
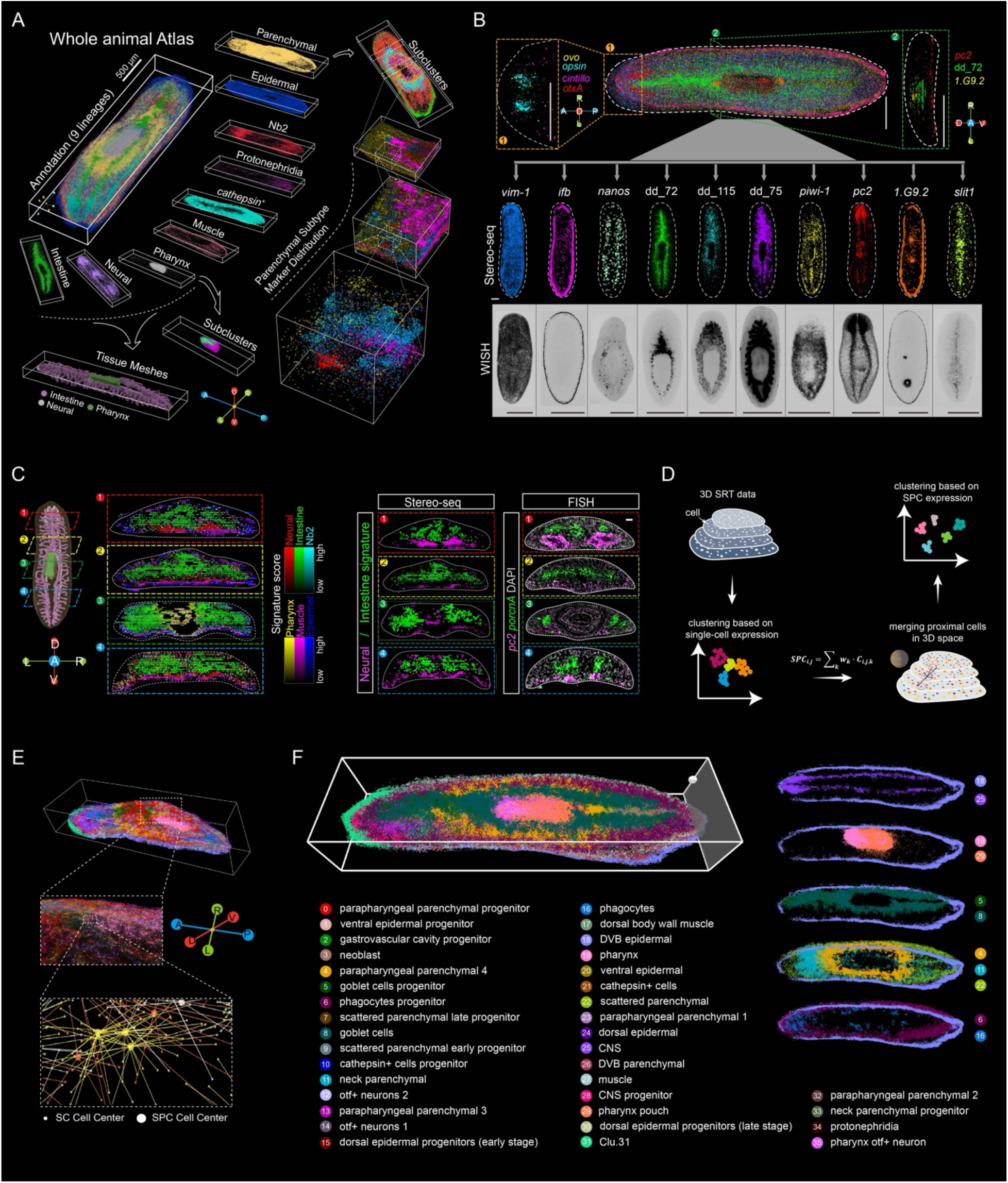
Virtual representation of the entire planarian at single-cell resolution. (A) 3D visualization of nine cell lineages in the whole planarian. The hierarchical 3D distributions highlight regionalized tissue meshes, cell lineages, subtypes, and marker distributions at multiple resolutions. (B) 3D spatial visualization of marker gene expression. Shown are the top view of visual system markers (top left), neural, intestine, and pharynx markers (pc2, dd_72, and G9.2) distribution on a virtual slice of a transverse tissue section (top right), and the top view of 10 lineage markers in the intact animal (middle). Corresponding WISH staining for these markers is shown at the bottom. n ≥ 6 animals show similar results; scale bars, 500 µm. (C) Spatial visualization of cell types including pharynx (yellow), muscle (magenta), epidermal (blue), neural (red), intestine (green), and *tgs1*^+^ Nb2 neoblasts (cyan), revealed by their expression and distribution in virtual transverse cross-sections of the 3D planarian. Colored boxes correspond to slicing positions shown in the schematic. Cells are color-coded by signature scores (left). Spatial visualization of neuronal and intestinal cells based on Stereo-seq data, with FISH staining for *pc2* (neural) and *porcnA* (intestinal) markers in corresponding cross-sections shown in the left schematic. DAPI, nuclei (gray). n ≥ 6 animals show similar results; scale bars, 50 µm. (D) Schematic flowchart of Spatial-Proximal Clustering (SPC). Putative single cells are grouped based on their Euclidean distance in 3D space, forming the SPC cells. These SPC cells are then re-clustered based on merged expression similarity and spatial proximity. (E) Schematic diagram illustrating how SPCs are organized in digital 3D space. Smaller dots represent single-cell (SC) centers, while larger dots represent SPC cells. Dots are connected to their spatially nearest neighbors and colored according to SPC clustering annotations. (F) 3D spatial visualization of cell types derived from spatial proximity-based clustering (SPC). Colors, numbers, and cluster annotations correspond to the identified clusters.

To benchmark the 3D model, we compared the expression patterns of known marker genes with whole mount in situ hybridization (WISH) data (Fig. 2B). The results demonstrated largely consistency between our model and the expected distribution of neoblasts (*piwi-1*) [26], intestinal cell subtypes (*dd_72, dd_115, dd_75*) [24], and epidermal cells (*vim-1, ifb*) [27, 28] along the A/P and D/V axes. Furthermore, our model captured the intricate structures of regionalized tissues, such as photoreceptors (*ovo, opsin*) [29] and testis primordia (*nanos*) [30] (Fig. 2B). The 3D atlas largely preserved spatial continuity along the mediolateral (M/L) and D/V axes, as illustrated by virtual transverse cross-sections of representative cell types (Fig. 2C, left). Although some distortion was observed along the D/V axis, relative cellular positioning remained consistent with known tissue distributions [24] (Fig. 2C, right). Overall, the reconstruction provides a faithful representation of planarian tissue organization across body axes.

Spatial transcriptomics enables the identification of spatially organized gene expression patterns and cell neighborhoods within intact tissues, providing a framework to relate molecular states to tissue architecture and putative functional domains. We hypothesized that incorporating spatial information into clustering analysis would enable identification of spatially coherent clusters, as tissue function often arises from cells with similar transcriptional profiles located in close proximity (Fig. 2D). To test this hypothesis, we developed a spatial proximity-based clustering (SPC) method, grouping cells based on transcriptional similarity and spatial proximity (Materials and Methods) (Fig. 2E). The application of SPC effectively delineated known tissue architectures and cell lineages, showcasing its ability to capture spatially organized transcriptional regulation (Fig. 2F and fig. S1B). The cell types identified by SPC were consistent with those identified through single-cell clustering, as validated by Pearson’s coefficients and spatial location consistency analysis (fig. S1, C and D, data S1). In comparison to unsupervised single-cell clustering algorithms designed for scRNA-seq, SPC revealed more intricate tissue structures, enabling the refinement of cell lineages, such as neural cells, neoblasts, and intestinal cells, into distinct subtypes (fig. S1E). For example, neoblasts were further subdivided into different progenitor subtypes, while muscle cells were classified into ventral and dorsal subtypes with unique spatial distributions (fig. S1E and data S1). To validate the spatial patterns identified by SPC, we compared the SPC cluster markers with data from fluorescence in situ hybridization (FISH) and Stereo-seq. The results demonstrated largely consistent spatial expression patterns, particularly for genes associated with intestinal, epidermal, neural, and polarity regulation (fig.S1F).

Further validation of the SPC-defined clusters was performed by comparing our 3D spatial data with published scRNA-seq atlases [24, 27, 31] (data S2). We integrated these datasets using the Harmony algorithm [32] and embedded them into the same UMAP dimensions. This integration resulted in a well-merged UMAP plot, confirming the consistency of our spatial data with published scRNA-seq atlases (fig. S2A). Manual annotation of the cell clusters based on marker genes from the integrated dataset revealed consistent lineage-level annotations across studies (fig. S2B), though minor discrepancies were observed at the subtype level (fig. S2C). While the cell types across datasets were largely consistent (fig. S2D), differences in cell proportions between scRNA-seq and spatial transcriptomics data were noted (fig. S3A), likely due to the sampling differences of spatial transcriptomics.

Through the alignment of spatial and integrated clustering annotations, we found that the SPC clusters provided additional spatial insights compared to previous cell type definitions from scRNA-seq atlases. For instance, *otf*^+^ neurons [33] were further refined into pharynx-specific and scatter-patterned subtypes, while epidermal cells were classified into distinct dorsal, ventral, and DVB subtypes (fig. S3B and data S2). The SPC clustering also identified an anterior-localized domain, Clu.31, which included spatially co-expressed epidermal, muscle, and neuron cells (Fig. 2F; fig. S3B, and data S2). Sub-clustering of neoblast cells by SPC exhibited less diversity compared to scRNA-seq clustering (fig. S3, C and D) [5], likely due to limitations in sequencing depth. Analysis of PCGs revealed distinct spatial distributions along the A/P axis (fig. S3E), consistent with prior reports [18]. By integrating transcriptional similarity with spatial proximity, SPC improves resolution of spatial domains relative to conventional clustering approaches. The resulting 3D spatial transcriptomic map provides a multi-scale view of gene expression and anatomical organization, illustrating the potential of this approach to connect molecular variation with tissue architecture at organismal scale.

### Spatial gene expression gradients define axis-patterned regulatory programs

To further investigate the exact mechanisms of morphogenesis in planarians, we leveraged spatial transcriptomics data to analyze gene distribution in intact planaria, focusing on genes that exhibited spatial variability across the body axes. We categorized these genes into 16 clusters along the A/P axis, 31 clusters along the M/L axis, and 73 clusters along the D/V axis (Fig. 3A). Furthermore, we mapped the remaining genes to these clusters, resulting in the identification of 12,449 genes that showed clear spatial gradients, termed spatially biased genes (SBGs) (data S3). These SBGs demonstrate well-defined expression patterns along the A/P, M/L, and D/V axes of the planarian body, providing insight into the spatial organization of gene expression.

**Fig. 3.**
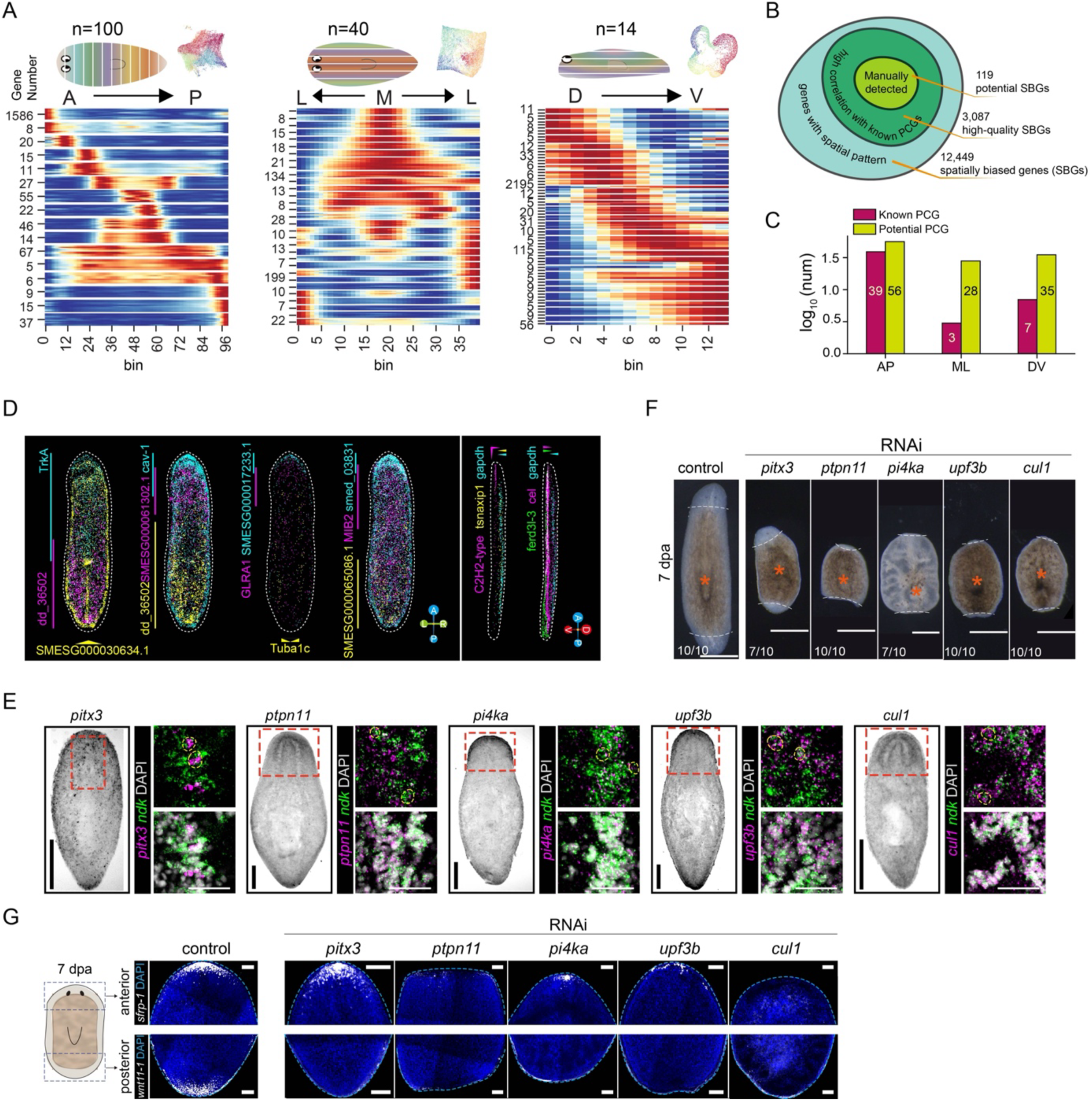
Generalized modeling and prediction of morphogenic gradient genes in homeostatic planarians. (A) Heatmaps showing clusters and smoothed average expression of spatial biased genes along the A/P, M/L, and D/V axes in the homeostatic planarian. Gene expression profiles are binned (top) and clustered using HDBSCAN. Genes that do not form clusters are re-categorized using logistic regression. (B) Schematic diagram illustrating the strategy used to identify PCG candidates, with the number of genes at each step indicated. (C) Bar plot depicting the number of known and newly predicted PCG candidates. (D) Spatial visualization of potential PCGs candidates in homeostatic animals using a 3D projection algorithm. (E) WISH showing the expression distribution of potential PCGs (left) and FISH showing co-expression of potential PCGs with *ndk* (right). Dotted circles indicate co-expressed cells. Scale bars: 500 µm (top), 50 µm (bottom); n ≥ 6 animals display similar results. (F) Representative phenotypes observed following knockdown of potential PCG candidates. Scale bars, 500 µm; n ≥ 6 animals show similar results. (G) FISH staining showing the expression and distribution of anterior and posterior markers following knockdown of potential PCGs. Scale bars, 100 µm; n ≥ 6 animals show similar results.

Understanding the interactions among these genes is critical to unraveling the complex processes underlying morphogenesis. We hypothesize that these spatial gradients might reflect potential interactions between PCGs and their downstream target genes. To identify genes that might function as PCGs, we employed a rigorous selection process. First, we excluded genes that were differentially expressed between cell types, as identified by SPC and single-cell clustering. This ensured that our focus was on genes exhibiting spatial gradients rather than those differentially expressed across cell types. We then calculated correlations between each gene and known PCGs, retaining genes with correlation values greater than 0.8 as high-confidence candidates. This process identified 3,087 high-quality SBGs (Fig. 3B and data S4).

To further refine the list of potential PCGs candidates, we used a projection algorithm to visualize the gradient patterns of these genes. Among the high-confidence SBGs, 119 genes exhibiting clear gradient patterns were selected as potential PCG candidates, expanding the known list of PCGs from 49 to 168 genes (Fig. 3, B and C; fig. S4A and data S5). Notably, newly identified PCG candidates such as *TrkA*, *bche,* and *dd_36502* showed expression patterns similar to established PCGs, including *fz5/8-4*, *sFRP-1*, and *sp5 [34, 35]*, particularly along the A/P axis (Fig. 3D and fig. S4B). We validated the gradient patterns of both known and potential PCGs through FISH assays (fig. S4C).

We next examined the distribution of both known and newly identified PCG candidates across the SPC clusters. Contrary to previous reports that PCGs are primarily expressed in muscle cells [9], our analysis revealed that PCGs were also widely distributed across multiple cell lineages, including neural and epidermal cells (fig. S4D and data S5). This suggests that non-muscle lineages may also carry positional information. Consistent with this, when we examined the expression of known PCGs in both our dataset and published scRNA-seq data, we found that these genes were not only expressed in muscle cells but also in various other cell types, such as neural and epidermal cells (fig. S4E).

To assess the functional relevance of selected candidate PCGs in polarity formation, we conducted a focused RNAi screen targeting genes co-expressed with the anterior pole marker, *ndk [12]*. Knockdown of genes such as *pitx3*, *ptpn11*, *pi4ka*, *upf3b*, and *cul1* (Fig. 3E) resulted in varying degrees of inhibition in blastema regeneration, leading to phenotypic defects such as one-eyed, eyeless, and tailless organisms (Fig. 3F). To further explore whether these defects were associated with disrupted polarity, we examined the spatial expression of anterior (*sFRP-1*) and posterior (*wnt11-1*) pole markers. Knockdown of the potential PCG candidates resulted in reduced expression of *wnt11-1* at the posterior pole, inhibiting tail formation or causing abnormal regeneration at the anterior pole, while the effects on *sFRP-1* expression varied (Fig. 3G). These results support a functional role for selected candidates in A/P axis remodeling and polarity regulation during regeneration.

In summary, our three-dimensional spatial transcriptomic dataset enables systematic classification of genes according to axis-related spatial patterns and expands the repertoire of candidate positional regulators. These findings provide a framework for dissecting positional control mechanisms underlying planarian regeneration.

### 3D analysis identifies intestinal cells in the neoblast niche

In the homeostatic state, neoblasts are dispersed throughout the planarian body, where they actively cycle and differentiate in response to local cues [36]. However, the cellular composition and potential functional roles of cells surrounding neoblasts remain incompletely characterized. Three-dimensional spatial transcriptomics provides an opportunity to profile tissue architecture without prior assumptions. To investigate how the local microenvironment may influence neoblast behavior, we quantified cellular components in close spatial proximity to neoblasts, focusing on their immediate neighbors. To minimize bias arising from regional heterogeneity, we defined a cumulative “meta-niche” that integrates both cell density and spatial distribution of neighboring cell types (Materials and Methods).

Our analysis revealed that most cell types were detectable within a 15 μm radius of neoblasts (fig. S5A). However, certain clusters, including dorsal epidermal progenitors and *cathepsin*⁺ cells, exhibited non-uniform spatial distributions relative to neoblasts (fig. S5A), suggesting regional patterning or context-dependent associations. Quantitative spatial proximity analysis further showed that gastrovascular cavity progenitors and phagocyte progenitors were the most frequent neighboring populations (Fig. 4A). This observation was supported by spatial visualization, which showed that neoblasts were often located near the terminal branches of the intestine (Fig. 4B). Enumeration of neoblast-adjacent cells confirmed enrichment of these intestinal-associated populations in close proximity to neoblasts (Fig. 4C), suggesting a potential role in shaping the local stem cell environment.

**Fig. 4.**
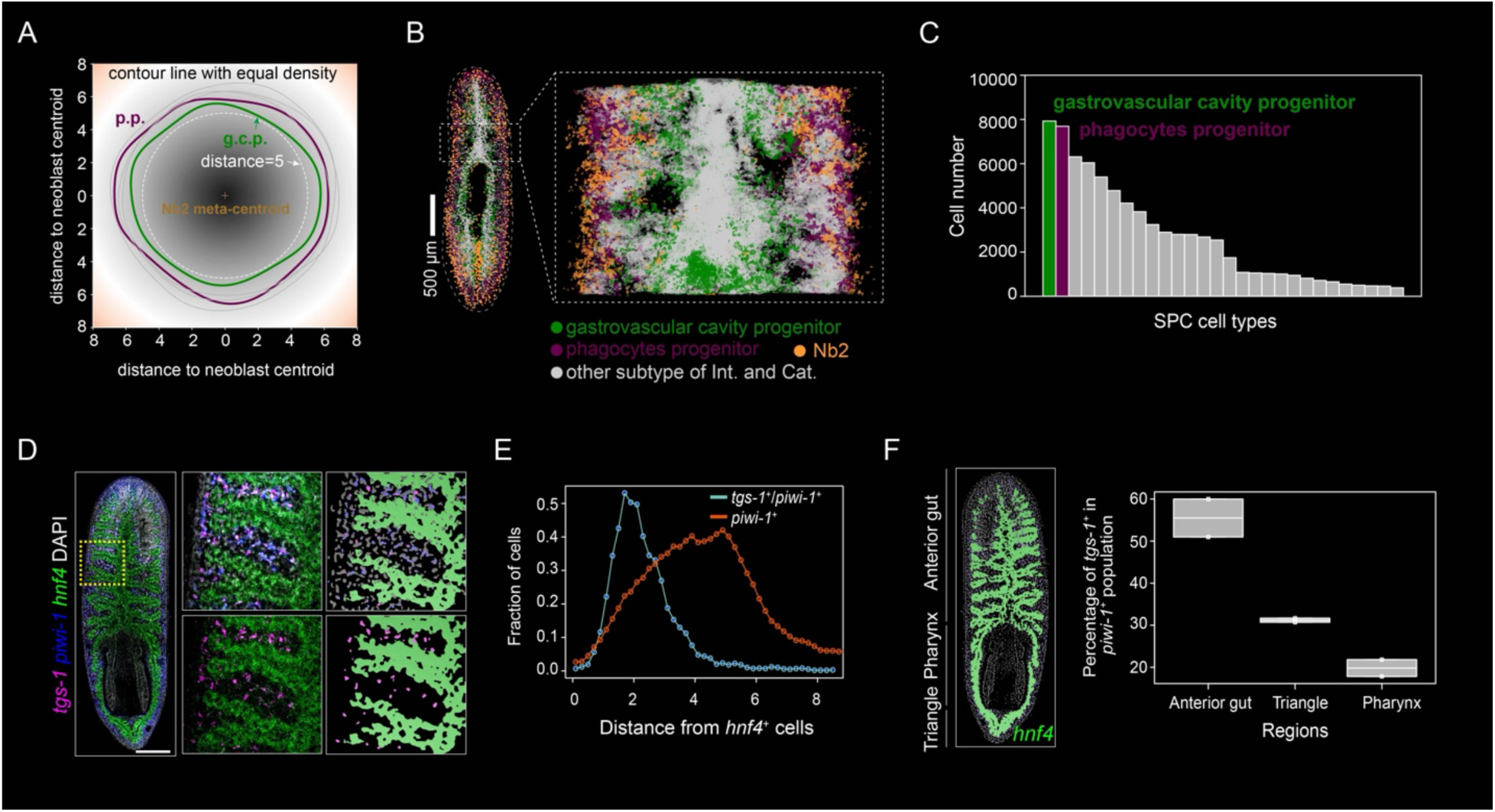
Detection of neoblast neighboring cells. (A) Contour map showing the top five SPC clusters closest to Nb2 cells. Kernel density estimation was performed for each SPC cell around Nb2 cells, with all Nb2 cell centers scaled to a single meta-centroid. The relative distances to the centered Nb2 were confined within a 15 µm radius. SPC clusters with contour line densities above 0.001 are displayed (Methods). Units for both x- and y-axes are in µm. p.p., phagocyte progenitor; g.c.p., gastrovascular cavity progenitor. (B) 3D spatial visualization showing the distribution of neoblasts and the two nearest SPC clusters. (C) Bar plot showing the number of cells in each SPC cluster within a 15 µm radius around Nb2 cells. The top five SPCs are highlighted, with colors representing different SPC types. (D) FISH staining of tgs-1 (magenta), piwi-1 (blue), and hnf4 (green) in homeostatic planarians (left). The pre-pharyngeal region is highlighted (middle), with 3D reconstruction using IMARIS (right). Scale bars, 500 µm. (E) Changes in the proportion of piwi-1+ or tgs-1+/piwi-1+ cells at varying distances from hnf4+ cells. (F) Left: 3D reconstruction of the whole animal in (D) using Imaris. Right: Boxplot illustrating the percentage of tgs-1+ cells in different regions surrounding the intestine.

To validate these findings, we performed FISH staining for neoblast markers (*piwi-1* and *tgs-1*) and an intestinal marker (*hnf4*) in homeostatic planarians (Fig. 4D). Three-dimensional FISH reconstruction enabled measurement of distances between neoblast subtypes and *hnf4*⁺ intestinal cells. Whole-animal analysis demonstrated that *tgs-1*⁺ neoblasts were positioned closer to *hnf4*⁺ cells than the broader *piwi-1*⁺ neoblast population (Fig. 4E), indicating preferential spatial association between *tgs-1*⁺ neoblasts (primarily Nb2 cells) and intestinal cells. In addition, *tgs-1*⁺ cells were enriched in the anterior gut region compared with triangular and pharyngeal regions (Fig. 4F). Together, these findings support a model in which intestinal cells, particularly those near terminal branches, represent prominent components of the *tgs-1*⁺ neoblast microenvironment. This spatial association is consistent with previous reports of functional interactions between intestinal and stem cell populations [37, 38].

## Discussion

This study establishes a semi-supervised workflow to address technical challenges associated with high-resolution three-dimensional reconstruction from spatial transcriptomics data, including complex tissue composition, blurred structural boundaries, and morphological distortions that complicate automated alignment [19, 20]. By integrating image registration, cell segmentation, clustering-based annotation, and serial section alignment, our approach enables three-dimensional reconstruction of complex biological structures at single-cell resolution. Applying this strategy to *Schmidtea mediterranea*, we generated a whole-animal 3D spatial transcriptomic atlas comprising nearly 900,000 segmented cells from an intact specimen. The resulting reconstruction captures the spatial organization of major cell types and gene expression gradients across body axes, consistent with prior scRNA-seq and in situ datasets. In addition, spatial proximity analysis identified intestinal cells as prominent neighboring components of neoblasts, and functional perturbation of selected positional gene candidates revealed roles in regeneration.

A central advance of this study is the integration of subcellular-resolution spatial transcriptomics with a semi-supervised three-dimensional reconstruction framework, enabling whole-organism 3D reconstruction at adult stage. Previous spatial transcriptomic analyses in planarians have primarily relied on two-dimensional sections or lower-resolution platforms [15, 39], limiting the ability to resolve individual cells and their spatial relationships across axes. In contrast, our approach combines high-resolution Stereo-seq data with multimodal image registration, adaptive cell segmentation, and serial alignment, enabling coherent reconstruction of a deformable and anatomically complex organism, revealing both consistent and unique features compared to previous definitions of planarian cell types [24, 27, 31]. Importantly, the incorporation of manual supervision at critical steps of alignment and segmentation allowed accurate reconstruction despite morphological distortions and variable tissue geometry. This hybrid strategy balances automation with biological constraint, providing a practical solution for reconstructing flexible soft-bodied organisms in three dimensions. By integrating spatial proximity information into clustering analysis, our framework further links transcriptional similarity with physical organization, facilitating the identification of functional domains that are not readily captured by scRNA-seq alone. Together, these features establish a scalable strategy for organism-scale 3D spatial transcriptomic reconstruction and extend the analytical resolution of spatial omics beyond static two-dimensional profiling.

Our analysis identified 119 candidate PCGs exhibiting clear spatial gradients across body axes. Notably, these candidates were not restricted to muscle cells, but were broadly distributed across multiple lineages, including neural and epidermal populations. Functional perturbation of selected genes resulted in regeneration defects, supporting their involvement in axial remodeling and polarity establishment. However, the contribution of additional candidates to pattern formation remains to be systematically evaluated. We further observed enrichment of PCG candidates within an anterior spatial domain (Clu.31) characterized by the co-expression of epidermal, muscle, and neuronal signatures in adult animals. Previous studies have emphasized muscle as a primary source of positional information during regeneration [40]. Our three-dimensional reconstruction refines this view by uncovering a broader and more complex distribution of spatially patterned genes across the body. These results suggest that positional information in adult planarians may be distributed across interacting cell lineages rather than restricted to a single tissue type.

Defining the stem cell microenvironment in highly regenerative organisms such as planarians is challenging due to the dispersed distribution and dynamic behavior of neoblasts [39]. To address this, we quantified the global composition of cells surrounding neoblasts under homeostatic conditions by constructing a cumulative “meta-niche” based on three-dimensional spatial measurements. This strategy provides a framework for systematically examining stem cell neighborhoods at organismal scale. Spatial proximity analysis revealed that intestinal cells, particularly those near branch terminals, are prominent neighboring components of neoblasts. The close spatial association between intestinal cells and *tgs1*⁺ neoblasts suggests potential physical and functional interactions, consistent with previous studies implicating the intestine in regulating neoblast activity and differentiation [37, 38, 41]. Further work will be required to identify the molecular signals mediating intestine–stem cell communication and to determine how these interactions contribute to tissue homeostasis and regeneration.

At present, our reconstruction is based on a single planarian specimen. Future analyses incorporating additional individuals will be important to evaluate the reproducibility and inter-individual variability of the observed spatial patterning. Despite this limitation, we examined dynamic changes in patterning across multiple regeneration time points, as described in an accompanying study (Han et al., submitted). Comparison with published scRNA-seq atlases [24, 27, 42], together with independent FISH validation, revealed substantial concordance and supports the overall reliability of our reconstruction. Although planarian stem cells (neoblasts) were readily identified and broadly consistent with prior datasets, finer neoblast subtypes reported in previous scRNA-seq studies [5] were not fully resolved. This likely reflects technical constraints inherent to spatial transcriptomics, including sequencing depth, sparse transcript capture, and partial mixing within spatial capture units, particularly given the small size of planarian cells. These considerations underscore the need for deeper sequencing and complementary high-coverage spatial transcriptomic approaches, as well as careful supervision during three-dimensional reconstruction, to further refine cellular resolution in complex organisms.

The relevance of this work extends beyond planarians. As basal bilaterians, planarians provide a useful system for examining conserved principles of cell type organization and positional gene regulation. The organism-scale 3D atlas presented here offers a reference for comparative analyses and illustrates an approach adaptable to additional model organisms. More broadly, this study contributes to the continued development of three-dimensional spatial transcriptomic strategies for interrogating complex tissues.

In conclusion, we present a semi-supervised framework for subcellular-resolution 3D reconstruction of whole organisms and provide a publicly accessible spatial atlas of adult planarians. This resource establishes a foundation for systematic investigation of cellular architecture, positional gene regulation, and regeneration at organismal scale.

## MATERIAL AND METHODS

### Planarian culture

Asexual *Schmidtea mediterranea* (strain CIW4) animals were maintained at 20°C in 1X Montjuic salts without antibiotics, as previously described [43]. The animals were transferred to static culture and starved for at least 7 days before all experiments.

### Gene cloning and RNA interference

Genes involved in this study were cloned from a CIW4 cDNA library and inserted into a pPR-T4P vector, as previously described [44]. The constructed plasmids, containing a T7 promoter, were used to induce the synthesis of dsRNA for RNA interference (RNAi). RNAi was performed to knock down specific genes, following established protocols [45, 46]. Briefly, dsRNA expression was induced in the bacterial strain HT115, and concentrated bacterial pellets were mixed with fresh beef liver paste in a 4:1 ratio. The mixture was stored at −80°C prior to feeding. dsRNA targeting EGFP was used as a control. Animals were fed every three days for a total of 1-2 RNAi feedings. On the third day after the final feeding, animals were amputated to collect regenerative samples.

### Whole mount *in situ* hybridizations

Whole mount in situ hybridizations was performed as previously described [47, 48]. Briefly, animals were incubated in 5% NAC in PBS to remove mucus, followed by fixation in 4% formaldehyde (FA) in PBSTx (0.5%) at room temperature for 1 hour. The animals were then bleached with formamide for 2 hours to minimize the influence of pigment. After a 2-hour pre-hybridization step, the corresponding RNA probe was added, and hybridization was carried out at 56°C for at least 16 hours. Following extensive washing, the probe was detected using an antibody, and signal amplification was achieved via the tyramide signal amplification system. Tissue clearing was performed by immersing the samples in ScaleA2 to reduce background interference during imaging [49]. Fluorescent images were captured using the Olympus SpinSR disk, and image processing was performed in Fiji/ImageJ. Robust gene expression was quantified as positive cells using CellProfiler.

### Tissue fixation, cryosection and ssDNA staining for Stereo-seq

Tissue fixation followed a previously described protocol with appropriate modifications [50]. Briefly, samples were euthanized in 0.66 M MgCl₂ for 1 minute and then fixed in Methacarn solution (6 mL methanol, 3 mL chloroform, 1 mL glacial acetic acid) for 10 minutes. After fixation, animals were rinsed three times with methanol, rehydrated in 50% methanol in PBS for 5 minutes, and subsequently dehydrated twice in 20% sucrose in PBS. The dehydrated tissues were embedded in pre-cooled OCT in a cassette, frozen on dry ice, and stored at −80°C until sectioning. Before sectioning, tissues were rewarmed at −20°C for 30 minutes. RNA was extracted from cryosections, and RNA quality was assessed using the Agilent 2100 Bioanalyzer.

Continuous 10 μm sections were prepared using a Leica CM1950 cryostat, and each section was mounted on a Stereo-seq chip. The slides were incubated at 37°C on a Thermocycler Adaptor for 3 minutes, followed by fixation with methanol at −20°C for 40 minutes. Single-stranded DNA (ssDNA) was stained using a nucleic acid dye (Thermo Fisher, Q10212) and visualized on a Leica DM6M microscope. Images were processed using the Leica Application Suite X software.

### Library construction and sequencing using Stereo-seq platform

The Stereo-seq library was constructed and sequenced following previously published methods [21]. Briefly, excess dye on the sections was washed with 100 μL of 0.1× SSC (Thermo, AM9770) containing 0.05 U/μL RNase inhibitor (NEB, M0314L). The slices were then incubated with 0.1% pepsin (Sigma, P7000) in 0.01 M HCl buffer (pH = 2) at 37°C for 18 minutes to permeabilize the tissues. Permeabilized mRNAs were captured on the Stereo-seq chip and reverse transcribed overnight at 42°C using the SuperScript II reverse transcription mix (Invitrogen, 18064-014), which contained 10 U/μL reverse transcriptase, 1 mM dNTPs, 1 M betaine solution, 7.5 mM MgCl₂, 5 mM DTT, 2 U/μL RNase inhibitor, 2.5 μM Stereo-seq template switch oligo, and 1× First-Strand buffer.

After in situ reverse transcription, residual tissue was removed, and the reverse transcription products were amplified using KAPA HiFi Hotstart Ready Mix (Roche, KK2602) with 0.8 μM cDNA-PCR primer. The PCR products were then fragmented using in-house Tn5 transposase at 55°C for 10 minutes. DNA fragments were ligated to adapters through PCR amplification using the same KAPA HiFi Hotstart Ready Mix (Roche, KK2602) with 0.8 μM cDNA-PCR primer. The purified libraries were sequenced on the MGI DNBSEQ-T1 sequencer (35 bp for Read1, 100 bp for Read2) following fragment distribution inspection. Following quality control of sequencing data, a spatial gene expression matrix at the subcellular scale was generated.

### Preprocessing of spatial transcriptomics data

Spatial transcriptomics data generated at subcellular resolution using the Stereo-seq platform required preprocessing before subsequent analysis. In the sequenced library, Read1 contained three components: coordinate identifiers (CIDs), molecular identifiers (MIDs), and polyT sequences. Read2 contained the corresponding cDNA sequence information captured at the same location. CID sequences were used to determine the x-y coordinates (715 nm resolution) of the cDNA, with a tolerance for 1-bp mismatches. The cDNA sequences were aligned to the genome of *S. mediterranea* (dd_Smes_G4), and only successfully paired reads were used for subsequent exon transcript analysis. The MID sequences provided unique molecular identifiers (UMIs) for transcripts, and PCR duplicates were removed using handleBam (https://github.com/BGIResearch/handleBam). Only read pairs with an MID quality score greater than 10 were retained for the construction of gene expression matrices with spatial information.

### MIRROR: Multimodal Image Registration by tRackline On-chip with enhanced Resolution

Gene spatial expression color-coded images were registered to their corresponding ssDNA staining images using a semi-supervised approach. The 2D gene expression images from Stereo-seq capture subcellular diversity (∼715 nm) [21], which distinguishes them from other in situ RNA capturing technologies that typically capture multiple cells within the same spatial spot. To improve precision and reliability, we developed the semi-supervised MIRROR pipeline for expression-ssDNA registration in a coarse-to-fine manner.

Two image files were prepared as input prior to registration. The carved track lines on the Stereo-seq chip surface provided precise marker information (∼2 μm width) for registration optimization. The ssDNA images captured less luminous intensity, while no RNA signals were present in the expression images. To enhance track line visibility, the greyscale image of ssDNA staining was first scaled based on camera specifications, and pixel brightness below a threshold was amplified. A median filter was applied to improve contrast (using the *prepareregistrationssda* function of GEM3D_toolkit). Meanwhile, gene expression matrices were converted to color-coded images, where each pixel (715 nm) represented a single spot, and the color magnitude reflected expression strength. Pixels at track lines were uniformly set to a value of 255 (using the *prepareheatmap* function of GEM3D_toolkit). Local contrast was further enhanced by integrating spatial expressions at different bin sizes.

The tissue morphology and track line grids were initially aligned using the TrakEM2 algorithm in Fiji. Isotropic rigid registration, including manual rotation, translation, scaling, and mirror flipping, was computed as an initial condition for subsequent fine registration. The morphological similarity was slightly affected by RNA random diffusion during fixation and permeabilization, which could reduce registration accuracy.

Subsequently, registration was refined based on the track lines. We developed an automated algorithm to associate track-line pairs and iteratively search for the global maxima of pixel overlap between paired track lines (using the *secondregistration* function of GEM3D_toolkit). Identifying the region of interest (ROI) was critical for restricting optimization to relevant regions using the ROI Manager in Fiji. This step improved computational efficiency and mitigated errors from periodic track-line patterns and image stitching. The improvement in registration quality was qualitatively assessed through visual comparison.

To further evaluate registration accuracy and determine whether it met the requirements for single-cell analysis, we manually identified approximately 2,000 pairwise landmarks between gene expression and ssDNA-staining images, uniformly covering relevant regions of six randomly selected 2D whole-body sections. These landmarks, defined by line-line intersections, were selected based on easily identifiable positions in both images. We calculated the Euclidean distance between these landmark pairs and generated basic statistics to quantify registration quality. The final registration error was found to be 1.47 ± 0.96 μm (2.05 ± 1.34 pixels), an improvement from the previous 2.52 ± 1.93 μm (3.52 ± 2.70 pixels) using TrakEM2. This average error is substantially smaller than typical cell sizes (∼10 μm).

All scripts used for 2D multimodal registration are available on GitHub: https://github.com/BGI-Qingdao/4D-BioReconX.

### Single-cell segmentation and assignment

We began by segmenting the nuclei from the ssDNA staining images and later transferred the cell boundaries to the aligned gene expression images. First, we reduced the background noise in the original ssDNA images using a global threshold. Local contrast was then enhanced using the CLAHE algorithm with a clip limit of 2 and a tile grid size of 8. Each ssDNA image was processed for five iterations to enhance the contrast between the nucleus and the background. Nuclei locations were detected using the Otsu adaptive threshold strategy, implemented through a customized CellProfiler pipeline *[22]*, without requiring manual annotation. The Mutex Watershed algorithm was then applied to automatically compute pixel affinities and predict the association of each pixel with a specific nucleus. RNA capture spots were assigned to individual cells based on whether they fell within the segmented cell boundaries.

For more complex nuclei segmentation, such as in crowded cell environments or out-of-focus regions, we used the Laplacian algorithm to calculate the blur value for each pixel by applying a sliding kernel within a 3-pixel window. Pixels with a blur value below 30 were flagged as undetectable and assigned to a bin partition (bin15, 10 µm in width and height), instead of being delineated by a boundary. After precise 2D registration of the gene expression images to the ssDNA-staining images, the boundaries detected for each cell or bin were mapped onto the spatial color-coded gene expression image. The resulting DNB-level gene expression matrices were aggregated into putative cells, ready for downstream analysis (gem2gemc function in https://github.com/BGI-Qingdao/4D-BioReconX).

### Dimensionality reduction and cell clustering

Since the cell volume was not fully segmented in 10-μm-thick sections for some cells, RNA or UMI counts per cell were normalized using the SCTransform function in Seurat (v4.0.2) *[51]* after removing cells with fewer than 50 UMIs. Reference-based integration was performed, along with reciprocal principal component analysis (RPCA), to mitigate the potential impact of batch effects.

PCA was then performed for dimensionality reduction, and the top 20 principal components were selected for unsupervised clustering. Graph-based Leiden community detection was applied in the PCA space with a resolution parameter of 0.8. Spatial information was not considered during clustering, as we aimed to evaluate clustering connectivity without coordinates, specifically to assess the accuracy and precision of in situ RNA capturing. The resulting clusters were identified using the FindAllMarkers function in Seurat with default parameters, which generated a list of differentially expressed genes (DEGs). Clusters were visualized using Uniform Manifold Approximation and Projection (UMAP) for dimensionality reduction.

### Cell type annotation at single-cell resolution

Segmented single cells were annotated to investigate genetic variations across cell types. We extracted the top 50 genes with the most significant P-values from the unsupervised clustering and queried them in publicly available databases (https://planosphere.stowers.org). Nine major cell lineages were previously defined: neoblasts, epidermis, muscle, neural, protonephridia, pharynx, intestine, cathepsin+, and parenchymal. Marker genes were further screened through gene set enrichment analysis, followed by manual annotation.

### SEAM: Serial sEction Alignment by anatomical and MRNA expression similarity

Cellular expression spots were masked based on their location in the pairwise ssDNA-staining image to exclude those outside the histological region, thus preparing the data for 3D registration. Overlapping or worn tissue regions and smudges were also masked to avoid introducing errors into the registration process. Single-cell-level annotation images were generated where the RGB values for cells from the same lineage across serial sections were consistent.

Serial annotation images were aligned based on similarities in morphology and annotation color codes, with each pair of neighboring images manually registered using the Fiji plugin TrakEM2. The z-axis space was set to 10 μm, corresponding to the thickness of the cryosections. As a result, all serial sections from the same individual were aligned within a unified 3D x-y-z coordinate system. A new set of coordinates for each spot was computed using the transformation matrices.

### Body straightening

The reconstructed model of the individual planarian was computationally straightened in 3D for quantitative comparisons based on morphological and molecular features.

The body straightening in the A/P-M/L (x, y) plane was based on the spatial distribution of PCGs. Initially, the conventional PCGs, *fz-5/8-3* (head-biased) and *wnt11-1* (tail-biased), which exhibit an expression gradient along the planarian A/P axis, were used to manually position the A/P poles. Next, we utilized the bilaterally symmetric expression pattern of the PCG *slit* to more precisely and smoothly define the A/P axis, in contrast to traditional anatomical-centerline-based straightening strategies. A cubic polynomial regression fitting function was applied to the x and y coordinates to generate the A/P curve, with the A/P poles constrained as fixed points. The coordinates of all 3D spots were then recomputed along the newly straightened A/P (x) axis, preserving their vertical distances from the axis as new M/L (y) coordinates.

For straightening in the A/P-D/V plane, we used the flatness of the reconstructed neural mesh model, rather than D/V PCGs, due to the blurry expression gradients of the latter. The cubic polynomial fitting function was again applied to the x and z coordinates of the neural mesh, with the A/P poles as the constraints. The region of the neural mesh near the pharynx was masked during the fitting process to avoid interference. The coordinates of all cells were then recomputed along the newly straightened A/P (x) axis, preserving their vertical distances from the axis as new D/V (z) coordinates.

### 3D spatial transcriptomics reconstruction of the whole animal body

The updated cell-gene expression matrices, aligned with 3D x-y-z coordinates and corresponding annotations, were preprocessed for 3D reconstruction and visualization. The spot cloud (approximately 1 million spots) was converted into 3D triangular meshes using 3D Slicer [52] via a manual method. Gaussian smoothing was then applied to the meshes using MeshLab [53].

### Spatial Proximity based Clustering (SPC)

Based on the assumption that functional units are composed of multiple cells rather than a single cell, and considering the dispersed distribution of neoblast cells in planarians, we employed a functional-spatial-proximity-based strategy to define hubs of cells that share similar states or functions. To begin, we re-clustered the putative single cells at a higher resolution of 1.6 for each individual, correcting for batch effects across different sections to ensure accurate clustering. This step allowed us to separate the interpretable cell clusters as distinctly as possible. Next, we grouped cells from the same cluster into spatially proximal subsets using linkage clustering, based on the Euclidean distance in 3D space. Each resulting group, referred to as a spatially proximal cell cluster, contained an average of 20 putative single cells.

For each spatially proximal cluster, we represented the meta-like cell by summing the raw counts to define its unique molecular identifier (UMI) and using the centroid location as its spatial representation. This approach significantly reduced computational complexity, decreasing the number of cells from millions to hundreds of thousands. Additionally, the non-adjacent aggregation of cells provided enriched data, capturing spatial heterogeneity due to the increased feature set.

### Spatial gradient gene profiling

We utilized a molecular coordinate system in 3D space, in addition to analyzing individual cells or SPC clusters, to quantitatively examine gene expression profiles, including temporal expression strength and spatial patterns. To simplify the analysis without compromising precision, we integrated gene expressions into 100, 40, and 14 bins along the A/P, M/L, and D/V axes, respectively, after straightening the body. This approach allowed us to better highlight spatial and temporal heterogeneity. For each bin, we calculated and averaged the SCT-transformed gene expressions. Along the A/P axis, the minimum x-coordinate was defined as the anterior pole, while the maximum was defined as the posterior pole. For the M/L and D/V axes, we excluded both anterior and posterior regions before binning to avoid the “banana problem” [54] associated with the 3D reconstruction of the curved planarian. The resulting average gene expressions were zero-normalized.

### Spatial pattern clustering

We began by identifying highly variable genes (HVGs) for each sample using Seurat, and compiled a list of previously characterized PCGs from published studies to serve as our gene set of interest. SCT counts were averaged per bin for each gene in this list. The expression data were then normalized by the total number of cells per bin, scaled to a range of −3 to 3, and smoothed using a Gaussian filter (sigma = 3). Genes with non-zero counts in at least five consecutive bins were retained for further analysis.

We applied a hierarchical density-based clustering algorithm to the filtered and normalized (scaled and smoothed) homeostatic gene expression dataset, grouping genes with similar expression and positional profiles along the body axis. This analysis revealed 16 distinct clusters, comprising a total of 5,726 genes along the A/P axis (data S3). Notably, 146 genes in clusters 1 and 2 were identified as anterior PCGs, consistent with previous studies [55], while 57 genes in clusters 14, 15, and 16 were enriched at the posterior end (data S3).

To optimize the clustering for each body axis, we fine-tuned the clustering parameters until the groups were visually distinct. For genes that were not initially clustered, we applied linear regression to map them to the group with the highest probability of membership.

### Identification of potential PCGs candidates

We calculated the correlation coefficient between each spatial gradient gene and known PCGs, retaining those with a correlation value above 0.8 as high-confidence candidates. Genes identified as differentially expressed in SPC and SC clusters were excluded, refining the candidate list to a set of high-quality, gradient-expressing genes. The expression patterns of each gene were visualized using a spatial projection algorithm, and those displaying distinct gradient profiles were manually selected as potential PCG candidates.

### Stem cell neighbor analysis in 3D spatial transcriptomics data

We defined the *meta-niche* as the composition of neighboring cells for each neoblast in 3D space, to assess the microenvironment of neoblasts under homeostatic conditions. Briefly, for a given region, we identified all cells within a 15 µm radius surrounding each neoblast and recorded their relative positions relative to the center of the neoblast. By scaling the evaluated neoblast location to a single central point, we calculated the number and density of each cell type in the vicinity of the neoblast. We ranked the most abundant and nearest cell types to evaluate their potential contribution to the microenvironment of the neoblast. For neighbor detection and calculation of nearest neighbor distances, we used the K-Nearest Neighbors (KNN) algorithm with a defined radius.

### Library construction and analysis for bulk RNA sequencing

Total RNA was purified using TRIzol from RNAi-treated animals. For each replicate, 2 μg of RNA was used to generate RNA libraries using the VAHTS Universal V8 RNA-seq Library Prep Kit (Vazyme, NR605). Libraries were sequenced on an Illumina NovaSeq S4 platform.

Raw reads were quality-checked and adapter-trimmed using Trim Galore (v0.6.7). Trimmed reads were mapped to the *S. mediterranea* genome (dd_smed_g4) using STAR (v2.7.10a). Read counts were quantified using featureCounts (v1.22.2), and genes with zero expression in all samples were excluded. Normalization and differential expression analysis were performed using the R package edgeR (v3.36.0). Genes with an FDR less than 0.05 and a logFoldChange greater than 1 or less than −1 were considered significantly differentially expressed.

### Web-based interactive database, PRISTA4D

To enhance the accessibility of the planarian cell atlas to the research community, we developed PRISTA4D (Planarian Regenerative Interactive Spatiotemporal Transcriptomic Atlas in Four Dimensions), a web-based, interactive database (https://www.bgiocean.com/planarian).

PRISTA4D is a free, open-source platform designed for data sharing and exploration in regenerative biology. It utilizes a customized, BSD-licensed Apache ECharts framework for the rapid development of web-based visualizations of the cell-gene atlas. This approach offers greater convenience for data and software maintenance compared to traditional local data viewers. Importantly, the database fetches metadata for each planarian along the regeneration timeline from a cloud-based data repository.

Written in JavaScript, PRISTA4D is cross-platform (compatible with Windows, macOS, and Linux) and supports modern browsers, including Internet Explorer 10+. The graphical user interface is built using the Element Vue UI toolkit (https://github.com/ElemeFE/element), while 3D dynamic visualizations are powered by the ECharts-WebGL extension pack. This enables seamless navigation of millions of data points in 3D, which can be freely browsed and rotated, thanks to the ClayGL WebGL and ZRender Canvas libraries.

PRISTA4D provides an accessible and extensible resource for exploring cell fate specification and differential gene expression in the regenerative planarian. To simplify data visualization for numerous cell clusters and thousands of genes, we have designed a user-friendly interface that allows users to select specific animals, view 3D anatomical models, identify interesting cells and genes, and adjust various parameters such as annotation versions and visual appearance.

## Acknowledgments

We thank L. Bolund and D. Little for the critical reading of the manuscript. This research was supported by the National Natural Science Foundation of China (32070828 to A.Z. and 32100514 to M.X.), the National Key R&D Program of China (2022YFC3400400), the National Key R&D Program of China (2020YFA0112502 and 2021YFA1100202 to A.Z.), the Strategic Priority Research Program of the Chinese Academy of Sciences (XDA16021300), the CAS Pioneer Hundred Talents Program (A.Z.), Shanghai Pujiang Program (20PJ1414600 to A.Z.), the Shanghai Science and Technology Committee (STCSM) (22ZR1468400 to A.Z.), Guangdong Genomics Data Center (2021B1212100001) and the Feng Foundation of Biomedical Research (A.Z.), Shenzhen Science and Technology Program (RCJC20221008092804002 to Y.G.).

## Author contributions

A.Z., X.X., G.F., H.L., K.H. and M.X. conceived and directed the study. A.Z., X.X., M.X. and H.L. supervised the work. M.Q.S., Y.C. and YR.L. performed the animal experiments. XW.L., W.G., JQ.W., W.W. and H.P. performed the Stereo-seq experiments. K.H., M.X., L.G., Y.L., Y.W. and Z.H. analyzed the data. L.G., Y.L. and T.Y. performed database construction. Q.L., L.Z. and X.M. assisted in the data analysis. R.Z., L.L., X.W., H.Z., X.S., S.L., W.Z., S.T.C, J.F., X.L., Y.G., J.W. and H.Y. gave relevant advice. A.Z., G.F., H.L., M.X. and K.H. wrote the manuscript with input from all authors.

## Data and materials availability

All materials used for stereo-seq are commercially available. All data generated in this study were deposited at CNGB Nucleotide Sequence Archive (accession code: STT0000028). Processed data can be interactively explored from our PRISTA4D database (https://db.cngb.org/stomics/prista4d; https://www.bgiocean.com/planarian). All original codes supporting the current study are hosted on GitHub (https://github.com/BGI-Qingdao/4D-BioReconX). The web-based interactive PRISTA4D database has also been made available (https://www.bgiocean.com/planarian). Any additional information required to reanalyze the data reported in this paper is available from the lead contact upon request.

---

**Table S1.**

Basic statistics of Stereo-seq data.

**Data S1.**

Marker genes of Seurat clusters.

**Data S2.**

Data and results from integration analysis with published scRNA-seq data.

**Data S3.**

Pattern clusters of PCGs along the body axes.

**Data S4.**

Correlation coefficients of spatial gradient genes with known PCGs.

**Data S5**

Cell type specificity of known and potential PCGs.

