## Supplementary Figures for "Cell Type Architecture and Positional Gene Gradients in an Adult Animal at Subcellular Resolution"

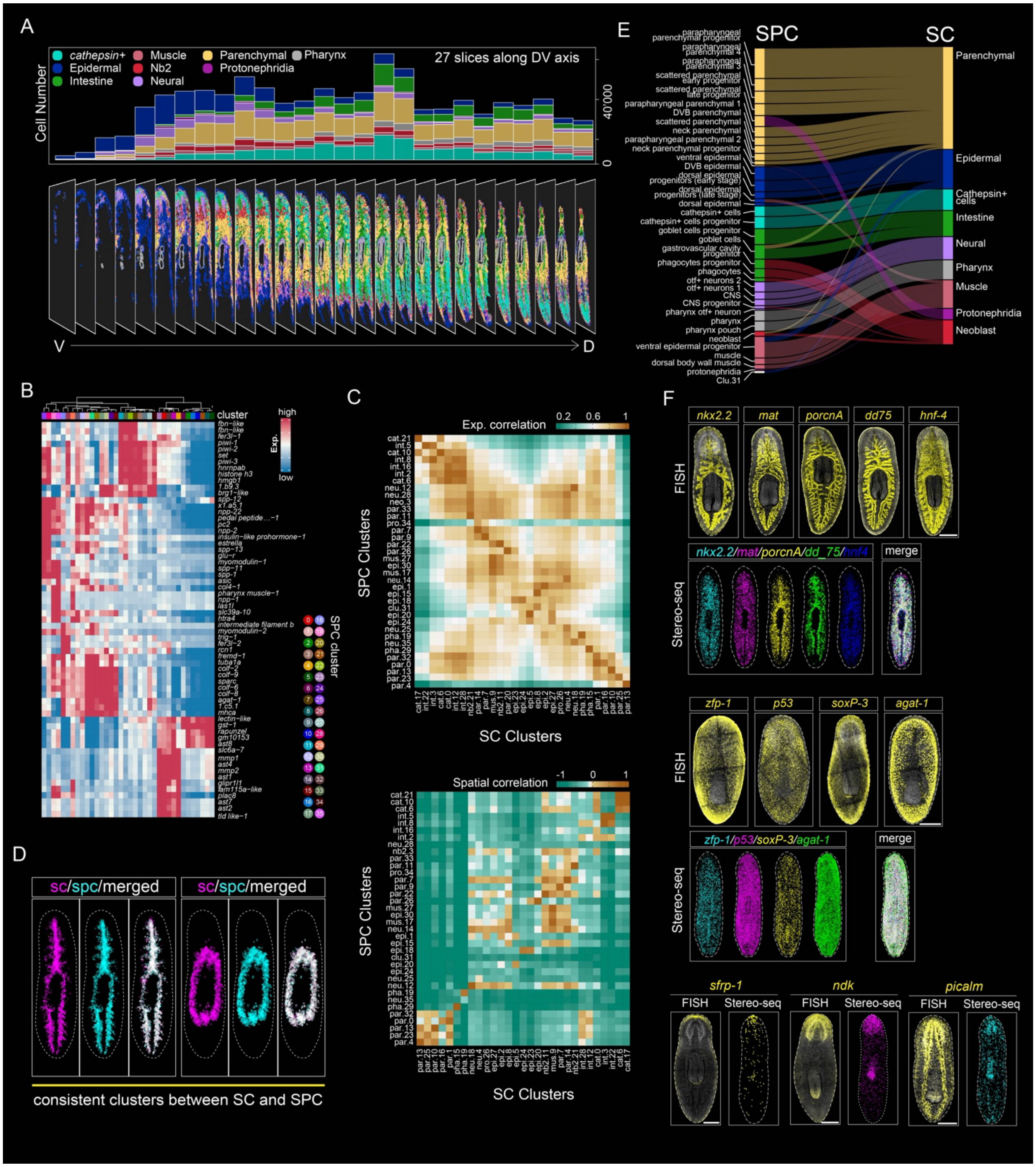


**Fig. S1. Sectioning information, clustering results, and validation of 3D reconstruction of a complete planarian.**

(A) Distribution of cell types across the 27 consecutively sectioned tissue samples used for 3D reconstruction. Top: Bar plots showing the distribution of nine cell lineages across each of the 27

Stereo-seq sections. Bottom: Spatial visualization of cell types from ventral (V) to dorsal (D), colored by their lineage annotations as shown in Fig. 2A. (B) Heatmaps illustrating the average expression levels of planarian cell lineage markers within SPC clusters. (C) Comparisons of single-cell (SC) and SPC clusters in terms of marker gene expression collinearity (left) and 3D spatial coordinate consensus (right). (D) Comparison of cell type visualization obtained from single cells (SC) or SPC, showing consistency in the parapharyngeal parenchymal (left) and intestinal cells (right). (E) Sankey plot illustrating the matching relationships between SPC and SC groups, with links colored according to lineage identity. (F) FISH staining showing the spatial expression and distribution of markers (top), with corresponding Stereo-seq data confirming consistent expression patterns of these markers (bottom). Scale bars, 500 µm; n ≥ 5 animals show similar results for FISH.

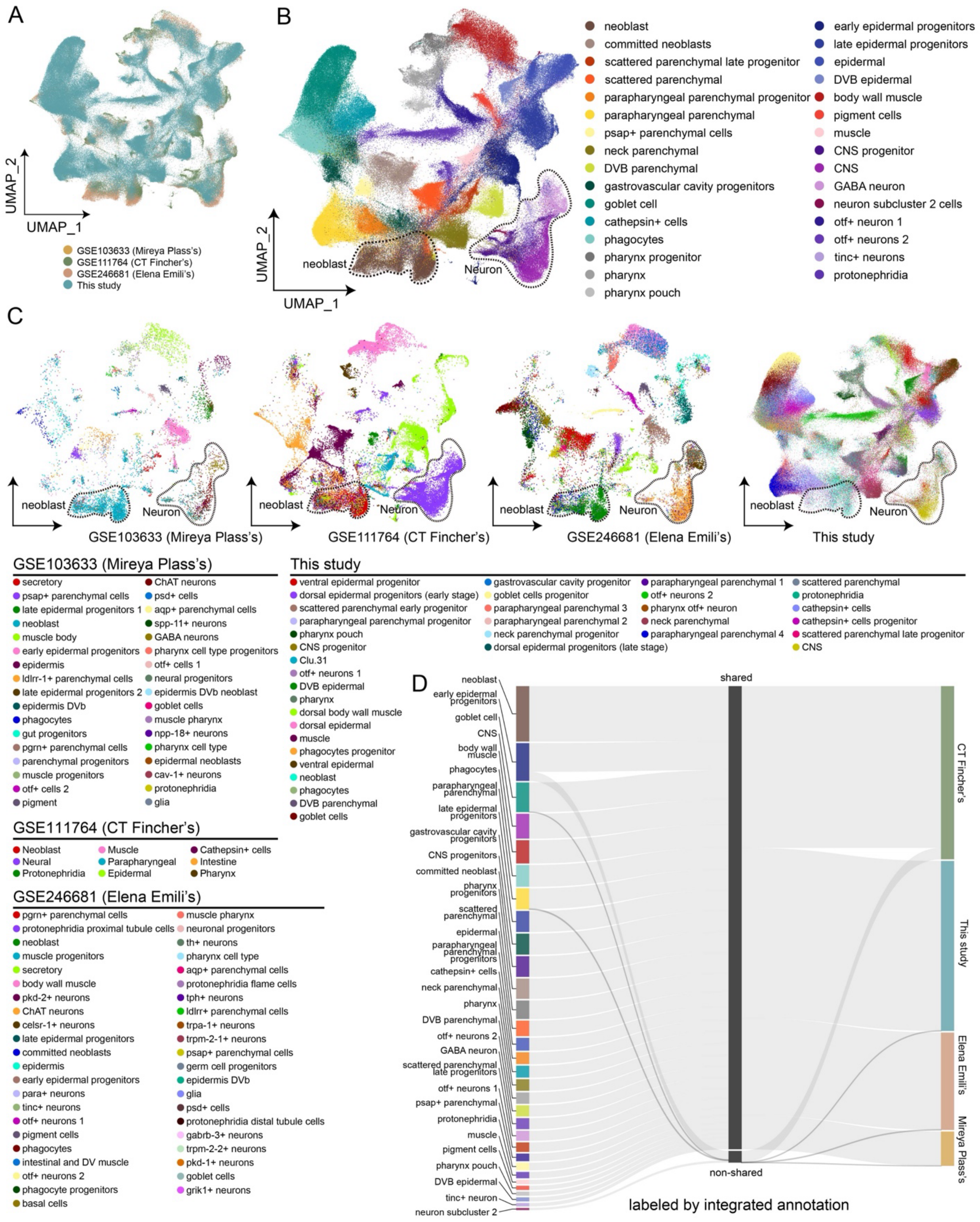
A
UMAP_2
UMAP_1
GSE103633 (Mireya Plass's)
GSE111764 (CT Fincher's)
GSE246681 (Elena Emili's)
This study
B
neoblast
Neuron
UMAP_2
UMAP_1
neoblast
committed neoblasts
scattered parenchymal late progenitor
scattered parenchymal
parapharyngeal parenchymal progenitor
parapharyngeal parenchymal
psap+ parenchymal cells
neck parenchymal
DVB parenchymal
gastrovascular cavity progenitors
goblet cell
cathepsin+ cells
phagocytes
pharynx progenitor
pharynx
pharynx pouch
early epidermal progenitors
late epidermal progenitors
epidermal
DVB epidermal
body wall muscle
pigment cells
muscle
CNS progenitor
CNS
GABA neuron
neuron subcluster 2 cells
otf+ neuron 1
otf+ neurons 2
tinc+ neurons
protonephridia
C
neoblast
Neuron
GSE103633 (Mireya Plass's)
GSE111764 (CT Fincher's)
GSE246681 (Elena Emili's)
This study
GSE103633 (Mireya Plass's)
secretory
psap+ parenchymal cells
late epidermal progenitors 1
neoblast
muscle body
early epidermal progenitors
epidermis
ldlrr-1+ parenchymal cells
late epidermal progenitors 2
epidermis DVb
phagocytes
gut progenitors
pgrn+ parenchymal cells
parenchymal progenitors
muscle progenitors
otf+ cells 2
pigment
ChAT neurons
psd+ cells
aqp+ parenchymal cells
spp-11+ neurons
GABA neurons
pharynx cell type progenitors
otf+ cells 1
neural progenitors
epidermis DVb neoblast
goblet cells
muscle pharynx
npp-18+ neurons
pharynx cell type
epidermal neoblasts
cav-1+ neurons
protonephridia
glia
GSE111764 (CT Fincher's)
Neoblast
Neural
Protonephridia
Muscle
Parapharyngeal
Epidermal
Cathepsin+ cells
Intestine
Pharynx
GSE246681 (Elena Emili's)
pgrn+ parenchymal cells
protonephridia proximal tubule cells
neoblast
muscle progenitors
secretory
body wall muscle
pkd-2+ neurons
ChAT neurons
celsr-1+ neurons
late epidermal progenitors
committed neoblasts
epidermis
early epidermal progenitors
para+ neurons
tinc+ neurons
otf+ neurons 1
pigment cells
phagocytes
intestinal and DV muscle
otf+ neurons 2
phagocyte progenitors
basal cells
muscle pharynx
neuronal progenitors
th+ neurons
pharynx cell type
aqp+ parenchymal cells
protonephridia flame cells
tph+ neurons
ldlrr+ parenchymal cells
trpa-1+ neurons
trpm-2-1+ neurons
psap+ parenchymal cells
germ cell progenitors
epidermis DVb
glia
psd+ cells
protonephridia distal tubule cells
gabrb-3+ neurons
trpm-2-2+ neurons
pkd-1+ neurons
goblet cells
grik1+ neurons
This study
ventral epidermal progenitor
dorsal epidermal progenitors (early stage)
scattered parenchymal early progenitor
parapharyngeal parenchymal progenitor
pharynx pouch
CNS progenitor
Clu.31
otf+ neurons 1
DVB epidermal
pharynx
dorsal body wall muscle
dorsal epidermal
muscle
phagocytes progenitor
ventral epidermal
neoblast
phagocytes
DVB parenchymal
goblet cells
gastrovascular cavity progenitor
goblet cells progenitor
parapharyngeal parenchymal 3
parapharyngeal parenchymal 2
neck parenchymal progenitor
dorsal epidermal progenitors (late stage)
parapharyngeal parenchymal 1
otf+ neurons 2
pharynx otf+ neuron
neck parenchymal
parapharyngeal parenchymal 4
scattered parenchymal
protonephridia
cathepsin+ cells
cathepsin+ cells progenitor
scattered parenchymal late progenitor
CNS
D
neoblast
early epidermal progenitors
goblet cell
CNS
body wall muscle
phagocytes
parapharyngeal parenchymal
late epidermal progenitors
gastrovascular cavity progenitors
CNS progenitors
committed neoblast
pharynx progenitors
scattered parenchymal
epidermal
parapharyngeal parenchymal progenitors
cathepsin+ cells
neck parenchymal
pharynx
DVB parenchymal
otf+ neurons 2
GABA neuron
scattered parenchymal late progenitors
otf+ neurons 1
psap+ parenchymal
protonephridia
muscle
pigment cells
pharynx pouch
DVB epidermal
tinc+ neuron
neuron subcluster 2
shared
non-shared
labeled by integrated annotation
CT Fincher's
This study
Elena Emili's
Mireya Plass's

**Fig. S2. Comparison of spatial transcriptomics data with published scRNA-seq datasets.**

(A) UMAP visualization of integrated spatial transcriptomics and single-cell RNA-seq data, with cells colored by their data GSE accession numbers. (B) UMAP visualization showing re-annotated cell types in the integrated datasets. (C) UMAP visualization comparing original cell types from the previously published studies indicated with GSE accession numbers with our SPC annotations. (D) Sankey plot illustrating the matching relationships between re-annotated cell types and each data accession.

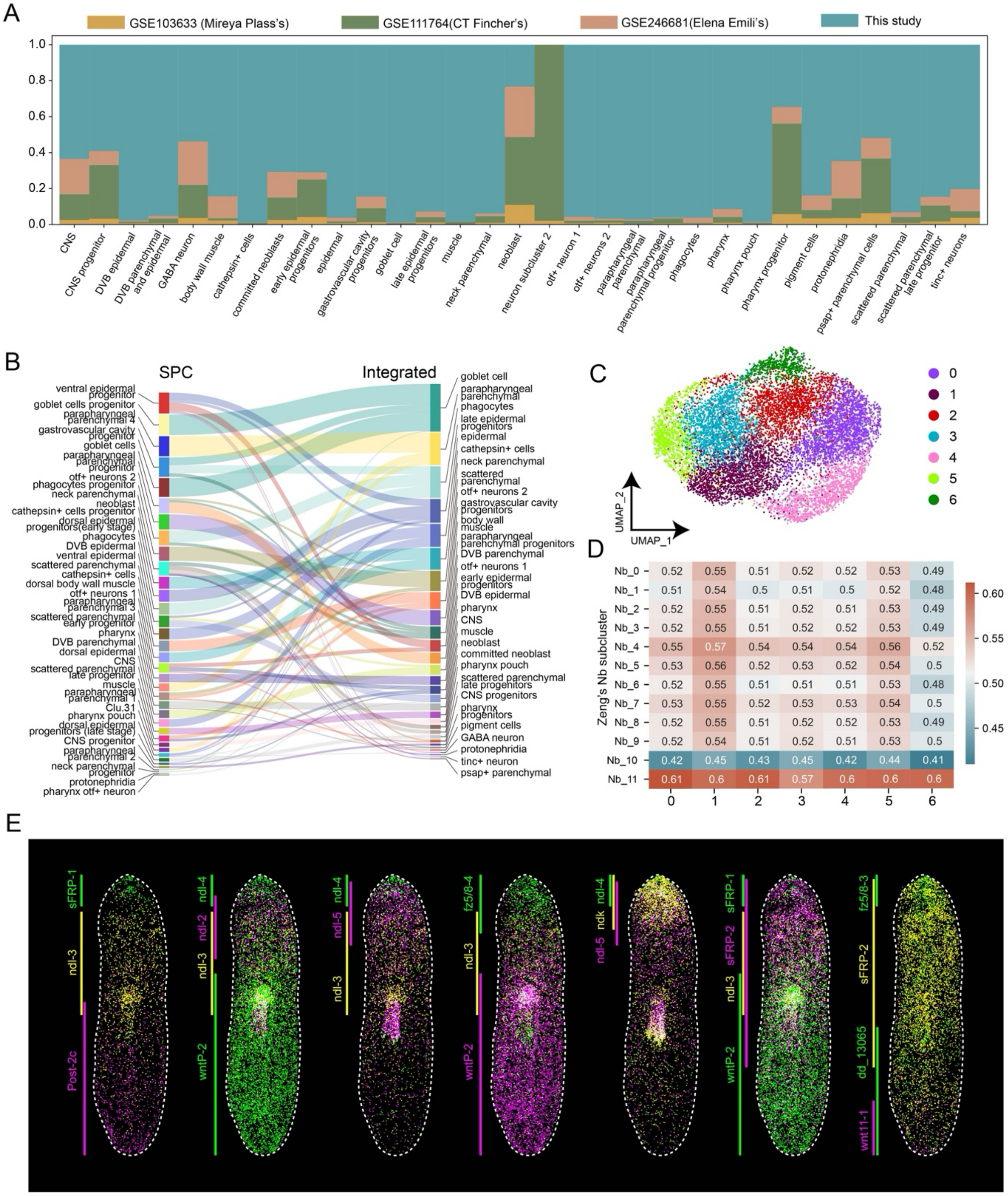


**Fig. S3. Characterization of stem cells and PCGs in spatial transcriptomics data.**

(A) Bar plot showing the percentage of cells from different data accessions for each re-annotated cell type. (B) Sankey plot illustrating the matching relationships between re-annotated cell types and spatial proximity-based clustering (SPC). (C) UMAP visualization of neoblast sub-clustering results. (D) Heatmap displaying the Spearman correlation between neoblast subclusters and previously published neoblast subtypes (Zeng et al., 2018). (E) 3D spatial projection visualizing known PCGs in uninjured planarians along the anterior-posterior axis.

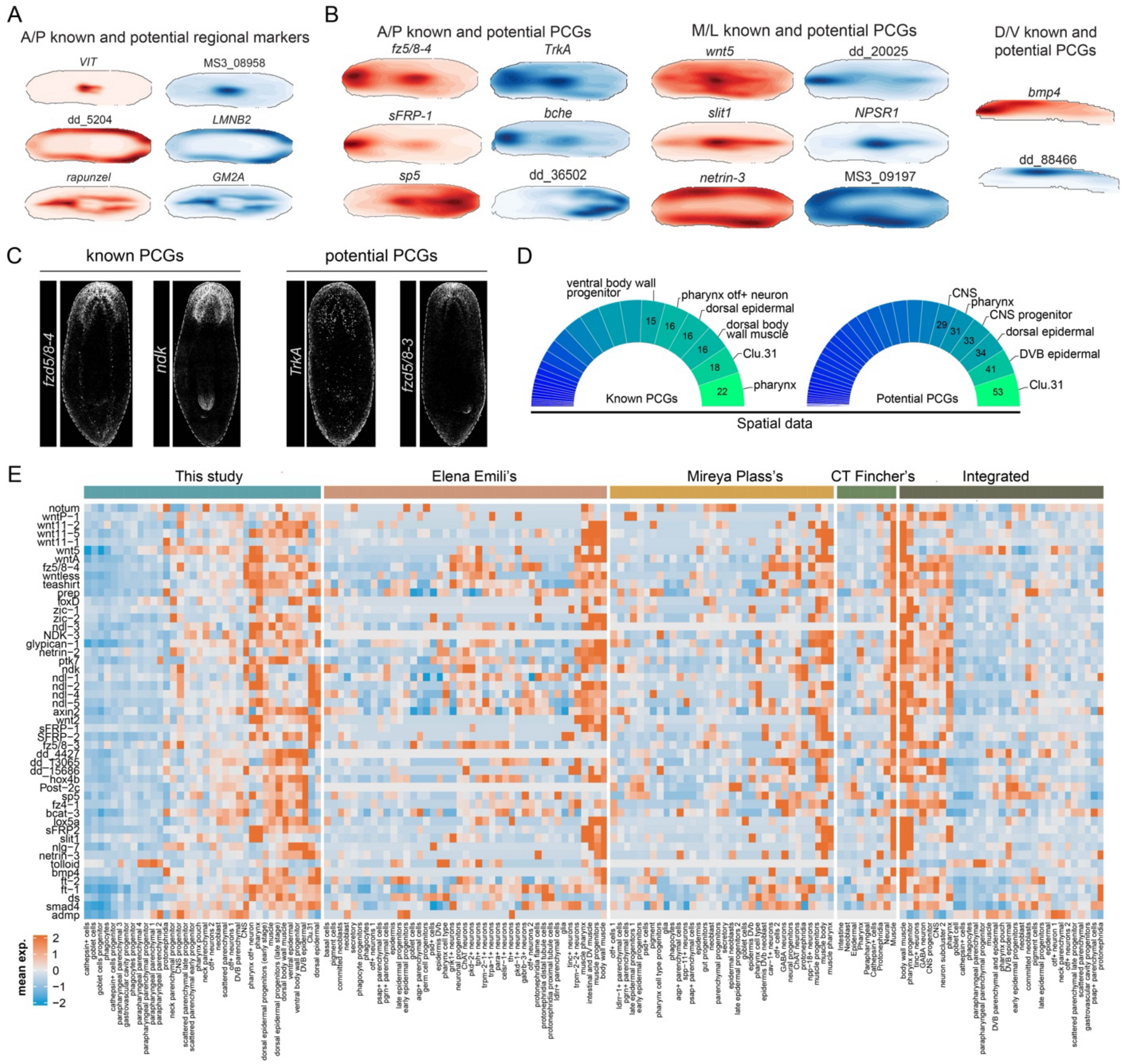


**Fig. S4. Analysis of expression of PCGs in spatial transcriptomics data.**

(A) Contour maps illustrating the gene expression profiles of marker genes relative to local histological structures. (B) Contour maps showing the predicted A/P, D/V, and M/L PCGs with global gradients. (C) FISH showing the expression of a subset of known and potential PCGs candidates in homeostatic animals. Scale bars, 500 μm; n ≥ 5 animals display similar results. (D) Pie chart illustrating the number of PCGs highly expressed in different spatial clusters. (E) Heatmap showing the average expression of known PCGs across different cell types in our dataset, as well as in published scRNA-seq and integrated datasets.

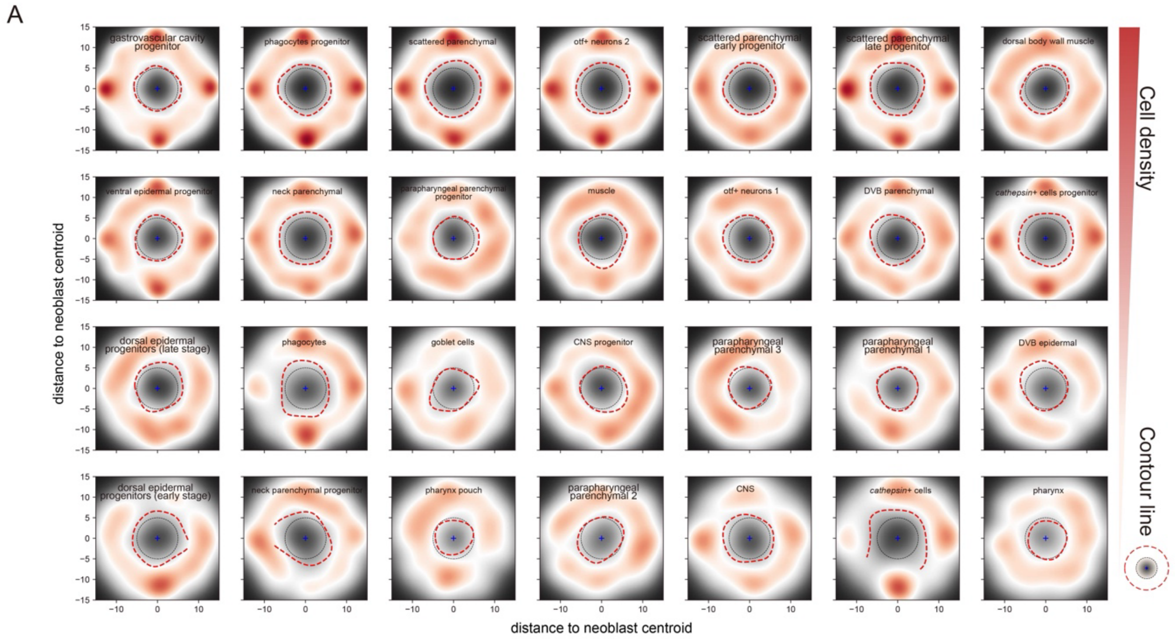


**Fig. S5. Analysis of neoblast neighboring components.**

(A) Kernel density plot showing the distribution of neighboring cells for each cell type to the neoblasts in the intact animals.

| | | |
|---|---|---|
| **Whole mount dataset** | section counts | 27 |
| | DNBs | 64,479,815 |
| | Body volume ($mm^3$) | 0.685 |
| | DV length (μm) | 355 |
| | ML length (μm) | 1,126 |
| | AP length (μm) | 4,324 |
| **single cell level** | segmented cell number | 1,258,917 |
| | cells with UMI greater than 50 | 893,703 |
| | average gene number per cell | 208 |
| | average UMI counts per cell | 357 |
| **SPC level** | SPC number | 49,651 |
| | average radius per SPC | 27.3128 |
| | average gene number per SPC | 1,876 |
| | average UMI counts per SPC | 6,177 |

**Table S1.**
Basic statistics of Stereo-seq data.

**Data S1. (separate file)**
Marker genes of Seurat clusters.

**Data S2. (separate file)**
Data and results from integration analysis with published scRNA-seq data.

**Data S3. (separate file)**
Pattern clusters of PCGs along the body axes.

**Data S4. (separate file)**
Correlation coefficients of spatial gradient genes with known PCGs.

**Data S5. (separate file)**
Cell type specificity of known and potential PCGs.
